# AI Models Excel at Orchestration but Falter at Biological Judgment: Findings from an Agentic Gene Annotation Study

**DOI:** 10.64898/2026.09.23.753765

**Authors:** Aritra Sinha

## Abstract

Large language model (LLM) agents are increasingly used both to direct biological analyses and to interpret their results. The core functions of agents-workflow control and biological adjudication-are often combined within the same agent and evaluated end-to-end, making it difficult to determine whether a model that is useful in one role is also reliable in the other. In this study, Genome Skeptic, a bacterial gene-annotation framework was developed in which all sequence-level measurements are generated by deterministic bioinformatics tools, while an AI controller can request follow-up analyses only from a predefined registry. In a prospectively locked, blinded cohort of 20 genomes balanced for *tet(A)/tet(B)* presence and absence, GPT-5.6 Sol acting as the workflow controller achieved the same accuracy as fixed and exhaustive strategies (18/20 correct) while requiring 32% and 47% fewer follow-up analyses, respectively. This was consistent with a 60-case prospective benchmark, in which markedly different evidence-acquisition policies produced different final evidence states but identical endpoint calls in all 60 cases. When the same Sol model was instead given final decision authority over the exact evidence state used by the deterministic adjudicator, accuracy fell from 18/20 to 11/20 (exact McNemar *P* = 0.016), specificity fell from 1.00 to 0.30, and the model corrected none of the deterministic errors. All nine endpoint disagreements shifted negative calls to unresolved rather than identifying missed positives. The same model that efficiently decided what to analyse next performed worse when asked to decide what the evidence meant. In this task, LLM performed well in orchestration and not biological adjudication.

## Introduction

Large language model (LLM) agents are moving from describing biological analyses to actively directing them. When connected to external software and databases, they can be very efficient to select tools, inspect outputs, request additional analyses and synthesize a final interpretation. Tool-augmented agents have shown benefits in chemistry (Bran et al., 2024), gene-set interpretation (Wang et al., 2025), protein-function annotation (Ponnapati et al., 2026) and single-cell omics analysis (Liu et al., 2026). In these systems, however, the same model often performs two fundamentally different functions. As a workflow controller, it decides what analysis should be performed next. As a biological adjudicator, it decides what the accumulated evidence ultimately supports.

These roles impose different requirements and have different failure consequences. A controller does not need to determine biological truth; it needs to choose an analysis that is likely to reduce the relevant uncertainty. A poor choice may waste computation or leave evidence incomplete, but its effect can be corrected by subsequent analyses or by a downstream decision layer. An adjudication error is different and more consequential in biological analyses; once the available evidence has been collected, an incorrect interpretation propagates directly to the reported biological conclusion. Thus, a model that is effective at selecting informative actions need not be equally reliable at converting evidence into a final call. Yet most agentic systems are evaluated end-to-end, so a single performance measure conflates evidence acquisition with evidence interpretation. ScienceAgentBench explicitly argued for evaluating individual scientific-agent capabilities before making claims about end-to-end automation (Chen et al., 2025), while systematic benchmarking in single-cell omics showed that planning, retrieval, reflection and execution contribute differently to overall agent performance (Liu et al., 2026).

A second question is whether effective workflow control requires a general-purpose LLM at all. To address this, Jev was included as a mechanistically distinct comparator. Jev is an RLCD-based, non-LLM typed decision model by Typesafe AI designed for constrained classification and decision tasks rather than free-form language generation (Almeida, 2026). Its inclusion allowed the contribution of general-purpose language reasoning to be separated from that of efficient structured decision-making. In the prospective experiment, Jev was therefore evaluated as a workflow controller alongside GPT-5.6 Sol, fixed control and exhaustive execution, providing a direct test of whether biological orchestration benefits specifically from an LLM or simply from an effective decision policy.

This understanding is particularly important in biology because recovering evidence and adjudicating its biological meaning are not equivalent computational problems. Biological evidence is often incomplete, redundant or context dependent, for e.g. sequence similarity may indicate common ancestry without establishing identical function; conserved domains may support membership in a broad family without resolving a specific paralogue; and several individually plausible signals may still support competing interpretations. Recent biological agents illustrate both the opportunity and the limitation. Wang et al. (2025) showed that grounding GeneAgent in curated biological databases improved gene-set interpretation and reduced unsupported statements relative to an ungrounded LLM. Ponnapati et al. (2026) similarly showed that tool access improved protein-function prediction. However, in both systems the language model still participated in synthesizing the final interpretation, so the contribution of tool orchestration could not be cleanly separated from the quality of biological judgement. Most directly, Zhang (2026) gave different agent configurations the same prompt and the same deterministic protein evidence and found only moderate reproducibility of the resulting biological relevance assignments, with high-confidence calls varying substantially between repeated runs. The agents reliably retrieved and organized evidence, but biological overinterpretation remained.

These observations motivate a more controlled question than whether an agent “works” overall-does the model add more value by deciding what evidence to acquire, or by deciding what that evidence means? To our knowledge, this has not been tested directly in biological annotation by placing the same model in each role while holding the underlying measurements, available tools and final evidence state constant.

Bacterial gene annotation provides a tractable system in which to make this separation. Primary measurements such as sequence identity, alignment coverage, profile scores and genomic coordinates can be generated reproducibly using established tools including BLAST+, MMseqs2, DIAMOND and HMMER (Camacho et al., 2009; Steinegger and Söding, 2017; Buchfink et al., 2021; Eddy, 2011). The harder problem is converting those measurements into a specific functional assignment when related families produce overlapping evidence. For e.g. the tetracycline-efflux determinants *tet(A)* and *tet(B)* provide such a case. Both belong to the major facilitator superfamily, a large and diverse group of membrane transporters, so identifying a genuine *tet(A)/tet(B)* determinant requires not merely detecting similarity to a transporter but discriminating the target family from related proteins and applying explicit criteria to the resulting evidence (Pao et al., 1998; Chopra and Roberts, 2001; Roberts, 2005).

In this study, Genomic Skeptic was developed as an evidence-constrained framework for bacterial gene annotation designed to separate evidence generation, workflow control and final biological adjudication into distinct computational layers. Sequence-level measurements were generated exclusively by deterministic bioinformatics tools, whereas the controller could select only from a predefined registry of follow-up analyses so that the contribution of adaptive decision-making could be evaluated independently of both measurement and endpoint assignment.

Within this architecture, OpenAI’s GPT-5.6 Sol was evaluated in two distinct roles. First, its ability to act as a workflow controller was tested by assessing whether analytical effort could be reduced without loss of accuracy while the final adjudicator was held fixed. Second, its performance as a biological adjudicator was examined by providing the model with a final evidence state identical to that assessed by an explicit rule-based decision layer. These effects were isolated using a prospective, blinded *tet(A)/tet(B)* cohort together with an exact evidence-state role-swap design. The objective was therefore to establish where within a biological workflow LLM-based decision-making adds value and where explicit, auditable decision authority remains more reliable.

## Methods

### System architecture

Genome Skeptic was designed to separate sequence measurement, selection of follow-up analyses and final biological adjudication. Sequence evidence was generated using BLAST+ 2.16.0, MMseqs2 18.8cc5c, DIAMOND 2.2.6 and HMMER 3.4 (Camacho et al., 2009; Steinegger and Söding, 2017; Buchfink et al., 2021; Eddy, 2011). Candidate *tet(A)/tet(B)* sequences were evaluated against a fixed target reference panel and predefined competing transporter families. Controllers could request only analyses from a predefined action registry and could not generate or modify sequence measurements. Final endpoint assignment was performed separately from workflow control (Fig. 1).

**Figure 1.**
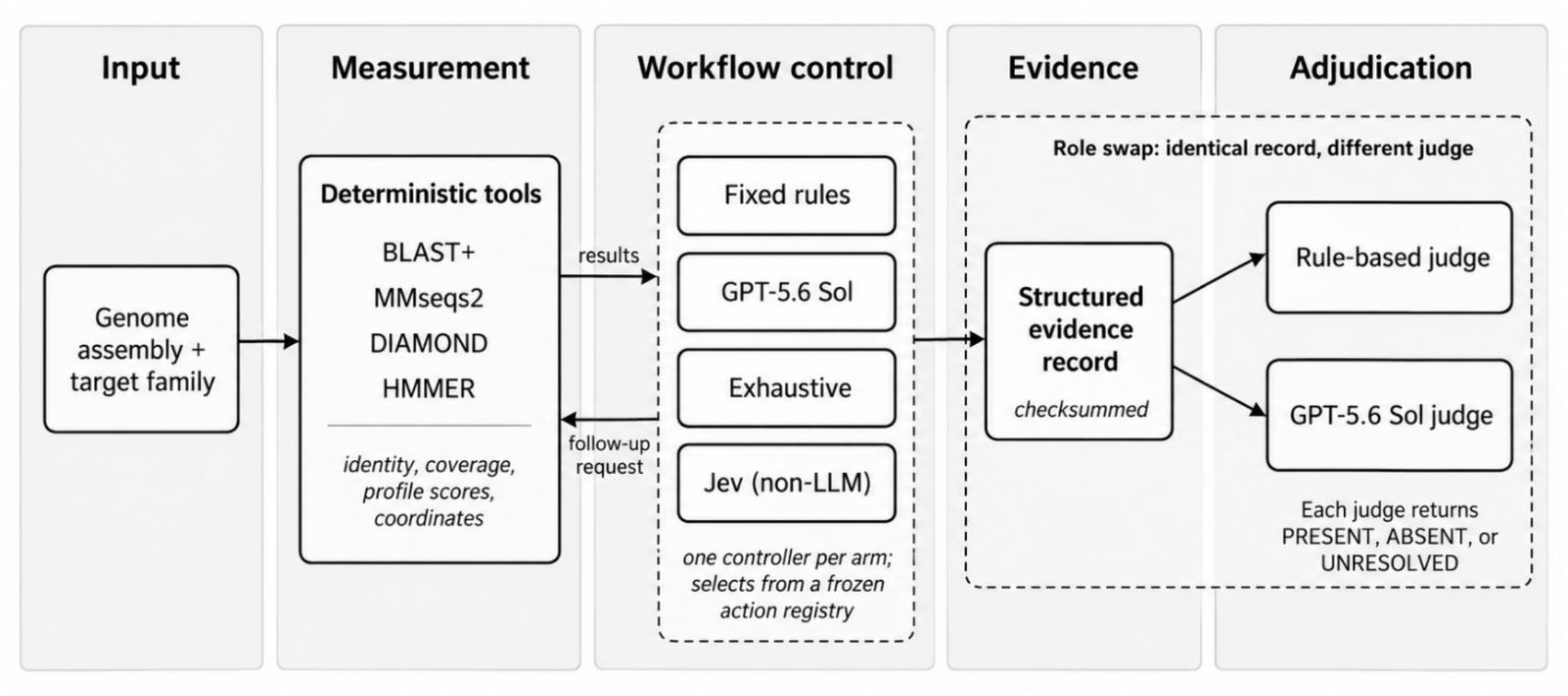
Experimental architecture of Genome Skeptic. Deterministic tools generate sequence-level measurements, while one controller per arm may request predefined follow-up analyses. The accumulated measurements form a checksummed evidence record. In the role-swap comparison, the identical evidence record is passed to either the rule-based judge or GPT-5.6 Sol, isolating the effect of final decision authority. Jev is a non-LLM typed decision model.

### Precursor study and system freeze

Before the prospective experiment, the framework was evaluated on 60 genome–target cases using fixed, Qwen3:4b and exhaustive controllers with the same rule-based adjudicator. All controllers shared the same eight errors, indicating that these failures arose from the decision layer rather than from follow-up analysis selection. Two adjudication defects affecting *tet(A)* family discrimination and *rpoB* profile coverage were corrected, after which the tools, reference panels, available actions and decision rules used for the prospective study were frozen.

Detailed analysis of the precursor errors and repairs is provided in the Supplementary Methods.

### Prospective cohort and truth assignment

The study protocol, truth criteria, case-selection procedure and experimental arms were fixed before prospective evaluation. Genomes used during development or as references were excluded. A balanced cohort of 20 genomes was selected, comprising 10 *tet(A)/tet(B)* positives and 10 negatives.

Truth was established using two distinct evidence routes independent of the systems under evaluation. The first compared recovered candidate ORFs against a frozen *tet(A)/tet(B)* reference panel and competing transporter families. The second evaluated the same candidates using target and competitor HMM profiles, with phylogenetic placement for borderline cases. Final truth required concordant positive or negative calls from both routes; discordant or near-threshold cases were classified as TRUTH_UNCERTAIN. All 20 selected genomes were concordant across the two routes.

The prediction pipeline had no access to truth labels during analysis. Outputs from all experimental arms were generated and locked before truth was joined for scoring. No case was replaced and no prediction was rerun after unblinding.

### Controller comparison and role-swap design

Four workflow-control strategies were evaluated using the same frozen rule-based adjudicator. Fixed control used a predefined action-selection policy. GPT-5.6 Sol was used as planner and critic, with at most one planner-selected and one critic-selected follow-up analysis per genome. An exhaustive controller executed every eligible follow-up analysis. Jev, a non-LLM typed decision model (Almeida, 2026), was evaluated under the same constrained control setting.

A separate role-swap condition isolated the effect of final decision authority. The final evidence record produced under GPT-5.6 Sol control was passed unchanged to GPT-5.6 Sol acting as the final adjudicator. No bioinformatics analysis was rerun, and the deterministic endpoint was withheld from the model. The evidence records supplied to the rule-based and LLM adjudicators were checksum-identical for all 20 genomes.

The Sol adjudicator was restricted to the locked evidence record and returned PRESENT, ABSENT or UNRESOLVED. Its structured output was required to reference existing evidence identifiers, and unsupported identifiers were rejected. No additional tool use or free-form chain-of-thought was permitted. The complete prompt and output schema are provided in the Supplementary Methods.

### Statistical analysis

The primary comparison was paired correctness between GPT-5.6 Sol control with rule-based adjudication and GPT-5.6 Sol control with LLM adjudication of the identical evidence state. Discordant outcomes were assessed using an exact two-sided McNemar test (McNemar, 1947). The paired difference in accuracy was estimated using case-resampling bootstrap confidence intervals. UNRESOLVED predictions were scored as incorrect according to the prespecified analysis plan, and proportions are reported with Wilson 95% confidence intervals (Brown et al., 2001).

Controller efficiency was summarized using the number of follow-up analyses, controller latency, total runtime and API cost. No inferential comparison of accuracy was performed among the four controller conditions because they produced identical final calls.

## Results

### Different controllers changed the evidence collected but not the final decision

In the 60-case precursor study, changing the controller altered the final evidence record in 30–45 of 60 cases but changed the final endpoint in 0 of 60 cases. All controllers shared the same eight errors. These errors were traced to two defects in the rule-based adjudication step rather than to evidence acquisition. The two rules were corrected and the system was frozen before the prospective experiment (Fig. 2).

**Figure 2.**
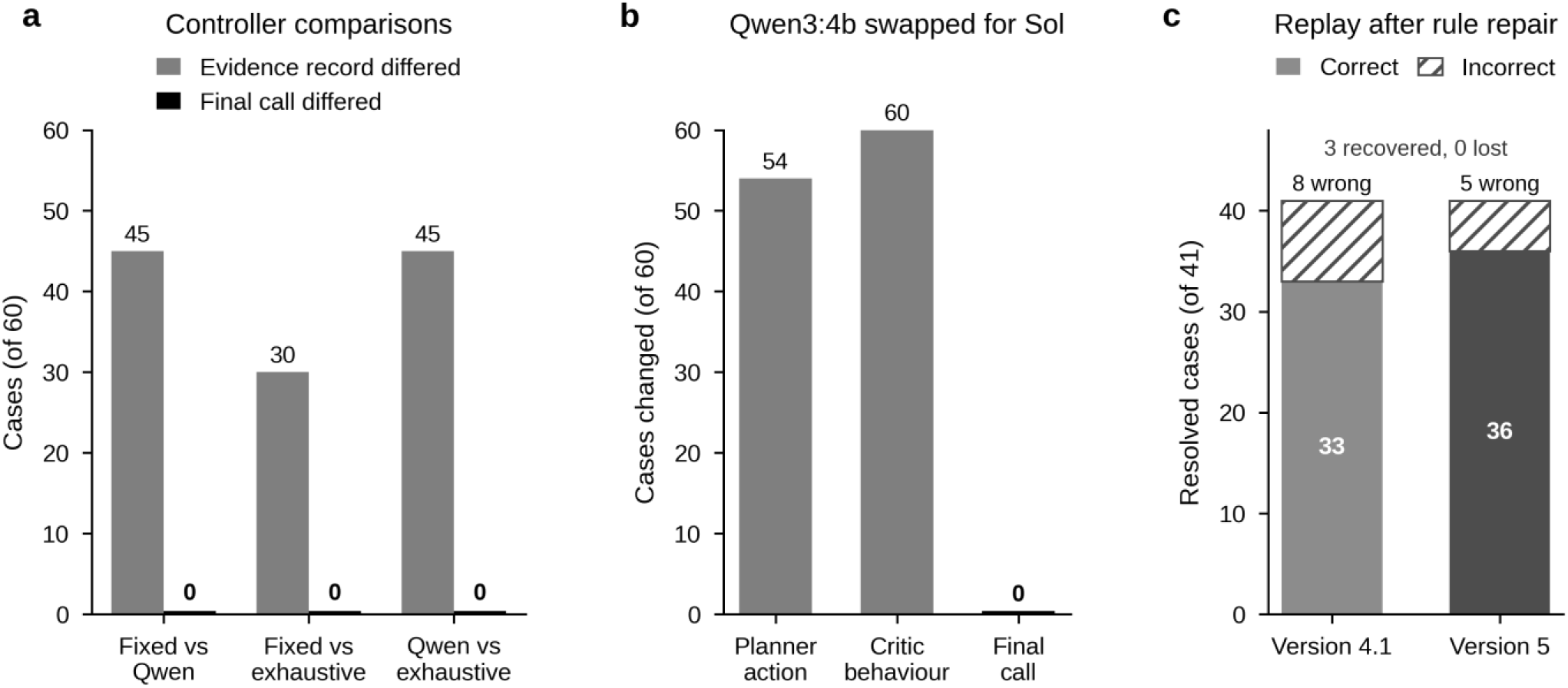
Precursor study. **a**, Pairwise comparisons of three controllers across 60 genome-target cases. Grey bars indicate cases in which final evidence records differed; black bars indicate cases in which the final call differed. **b**, Effect of replacing the Qwen3:4b controller with GPT-5.6 Sol, performed post hoc. **c**, Retrospective replay of the 41 truth-resolved cases before and after correction of two decision rules. Hatched segments denote incorrect calls.

This precursor result established that controller performance could only be evaluated meaningfully once the final adjudication layer was held constant.

### GPT-5.6 Sol reduced follow-up analysis without changing accuracy

In the prospective 20-genome cohort, fixed control, GPT-5.6 Sol control, exhaustive control and Jev control produced identical endpoint calls when evaluated by the same rule-based adjudicator. Each strategy classified 18 of 20 genomes correctly, with sensitivity of 8/10 and specificity of 10/10 (Table 1).

**Table 1.** Classification performance on the 20 prospective genomes. Values in brackets are Wilson 95% confidence intervals for accuracy. UNRESOLVED calls were scored as incorrect, as pre-specified.

| Arm | Controller | Judge | Follow-ups | Correct [95% CI] | Sensitivity | Specificity | Balance d accuracy | Unresolved |
| --- | --- | --- | --- | --- | --- | --- | --- | --- |
| A | Fixed rules | Rule-based | 57 | 18/20<br>[0.70, 0.97] | 8/10 | 10/10 | 0.90 | 0 |
| B | GPT-5.6 Sol | Rule-based | 39 | 18/20<br>[0.70, 0.97] | 8/10 | 10/10 | 0.90 | 0 |
| C | Exhaustive | Rule-based | 74 | 18/20<br>[0.70, 0.97] | 8/10 | 10/10 | 0.90 | 0 |
| F | Jev, non-LLM | Rule-based | 39 | 18/20<br>[0.70, 0.97] | 8/10 | 10/10 | 0.90 | 0 |
| D | GPT-5.6 Sol | GPT-5.6 Sol | 39† | 11/20<br>[0.34, 0.74] | 8/10 | 3/10 | 0.55 | 9 |

† Arm D adjudicated the evidence records generated in arm B; no analysis was re-executed.

The controllers differed substantially in analytical effort. GPT-5.6 Sol required 39 follow-up analyses, compared with 57 under fixed control and 74 under exhaustive control, corresponding to reductions of 31.6% and 47.3%, respectively. Despite identical final calls, the evidence record generated by Sol differed from that generated by fixed control in 12 of 20 genomes (Fig. 3).

**Figure 3.**
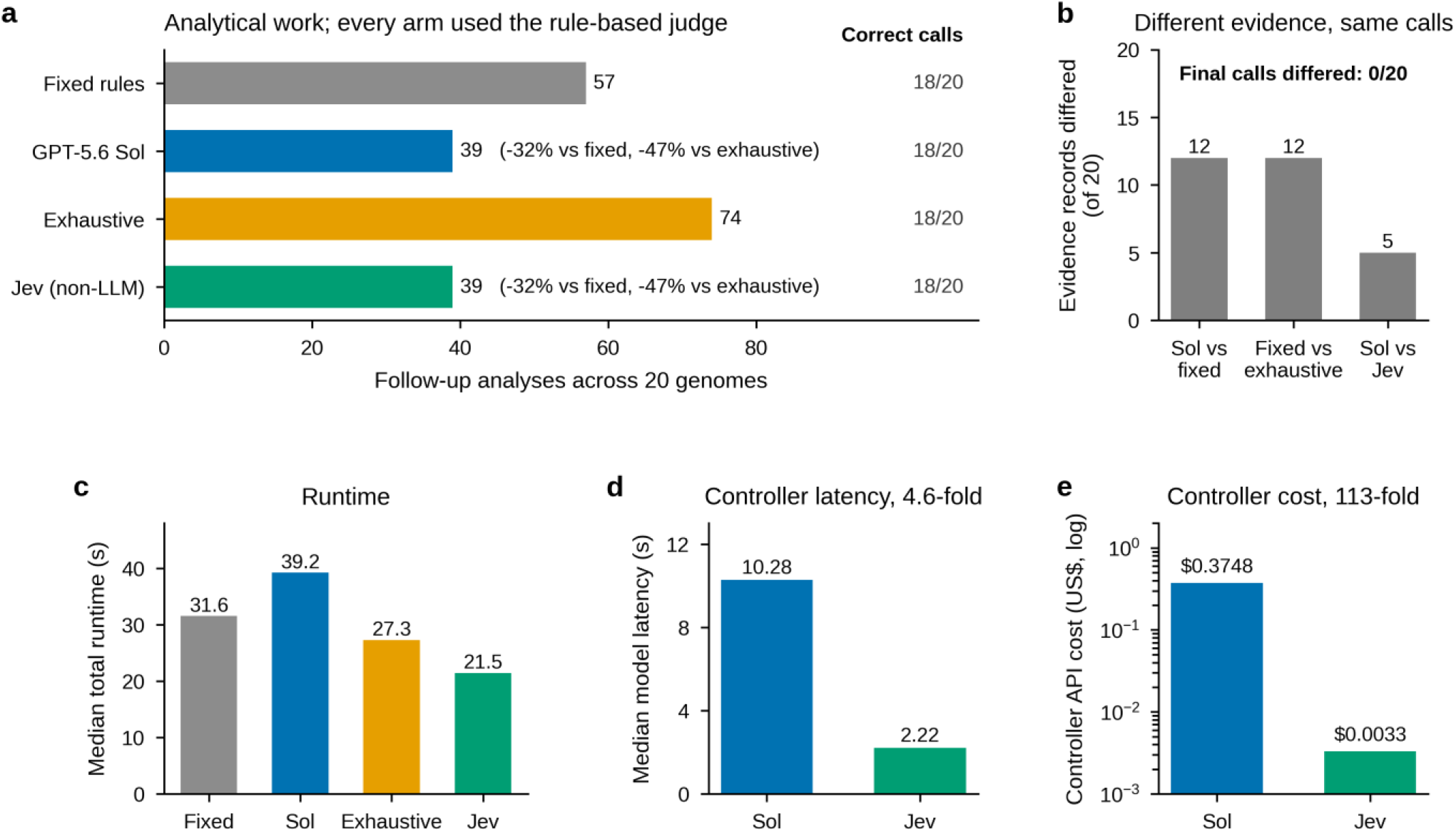
As a workflow controller, the LLM reduced analytical effort without altering classification. All arms used the rule-based judge (n = 20 genomes). **a**, Total follow-up analyses per controller, with correct calls shown at right. **b**, Pairwise differences in final evidence records; final calls were identical in every comparison. **c**, Median total runtime. **d**, Median controller-model latency. **e**, Controller API cost, on a logarithmic scale.

The reduction in analysis did not translate into lower runtime because of model inference latency. Jev, a non-LLM typed decision model, also required 39 follow-up analyses and produced the same 18/20 endpoint calls, while operating at 4.6-fold lower controller latency and 113-fold lower controller cost than Sol.

### GPT-5.6 Sol performed worse when given final decision authority

The role-swap experiment isolated the effect of adjudication. The final evidence records generated under Sol control were passed unchanged to either the rule-based adjudicator or GPT-5.6 Sol. Checksums confirmed identical evidence inputs for all 20 genomes, and no analysis was re-executed.

With the rule-based adjudicator, 18 of 20 genomes were classified correctly. When GPT-5.6 Sol adjudicated the same evidence, accuracy decreased to 11 of 20. Sensitivity remained 8/10, whereas specificity decreased from 10/10 to 3/10. The paired difference in accuracy was −0.35 (95% CI, −0.55 to −0.15), with an exact two-sided McNemar test of P = 0.016 (Fig. 4).

**Figure 4.**
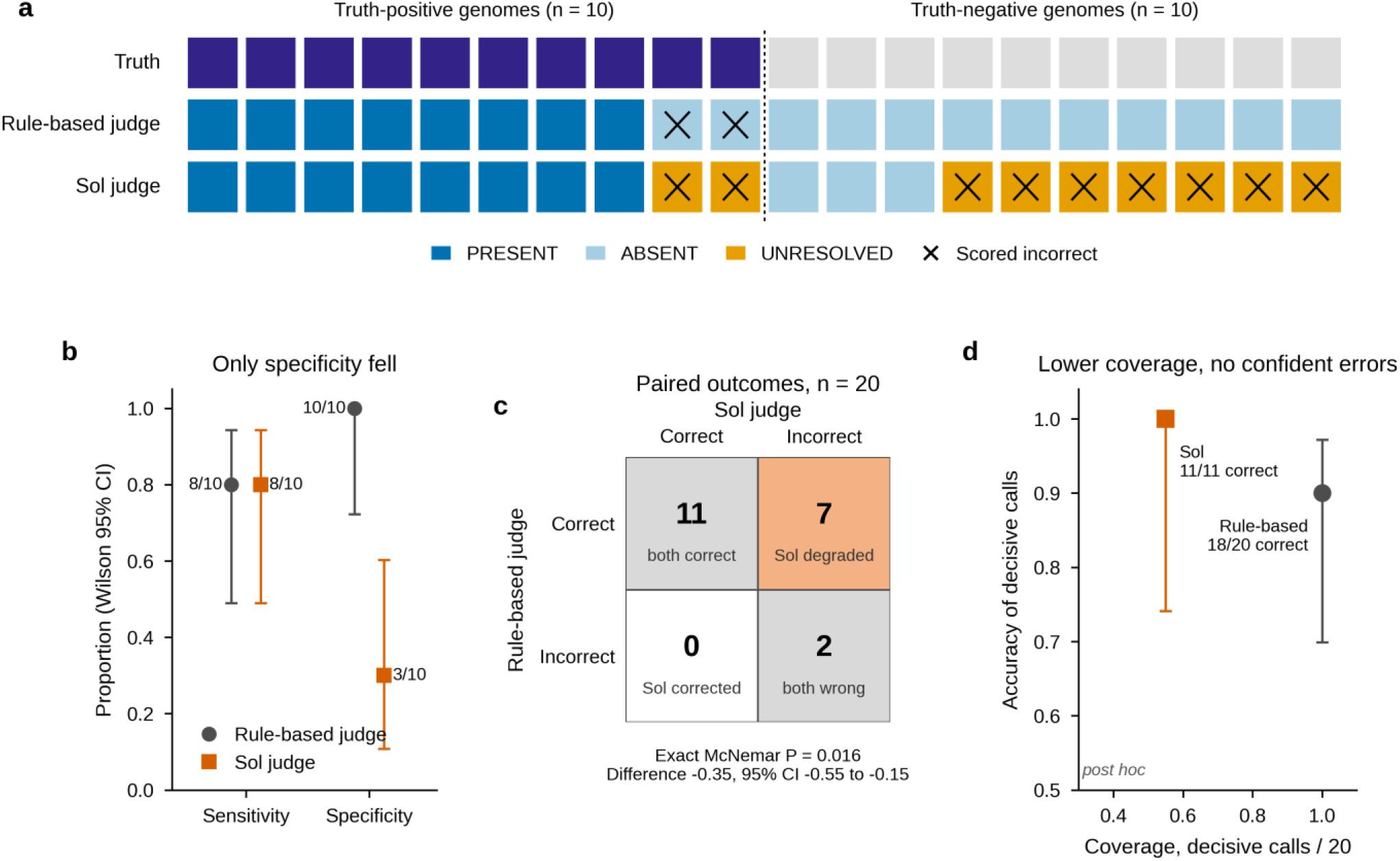
As a final adjudicator of identical evidence, the LLM was less reliable than explicit rules. **a**, Calls for each of the 20 prospective genomes, grouped by ground truth and ordered by outcome. Crosses mark calls scored as incorrect; UNRESOLVED was scored as incorrect, as pre-specified. **b**, Sensitivity and specificity with Wilson 95% confidence intervals. **c**, Paired outcomes for the two judges; only the discordant cells contribute to the exact McNemar test. **d**, Coverage, the fraction of genomes receiving a decisive call, against the accuracy of decisive calls, with Wilson 95% confidence intervals (post hoc analysis).

**Figure 5.**
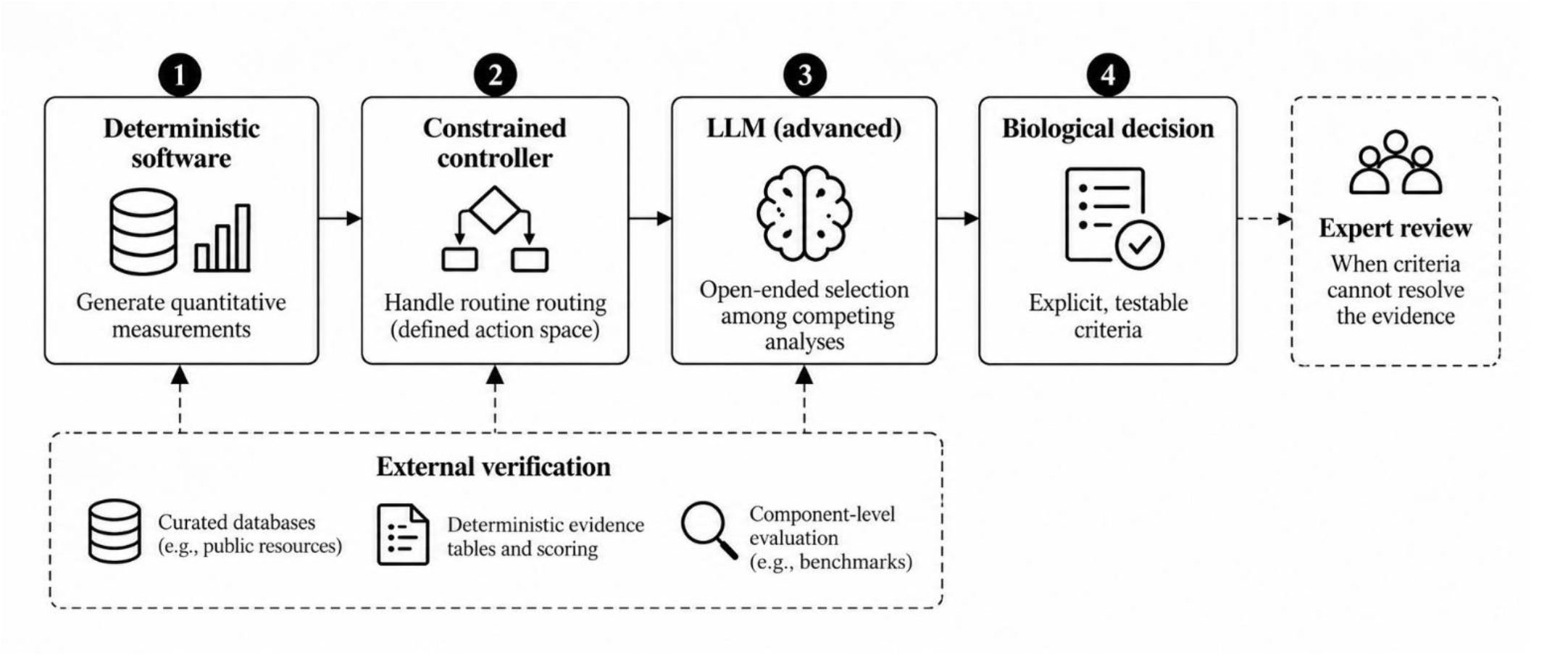
A proposed tiered architecture for scientific agents. This design assigns each component to the part of the workflow it handled most reliably in this study: deterministic systems for measurement and final judgement, lightweight controllers for efficient routing, and LLMs for higher-level orchestration.

There were nine endpoint disagreements between the two adjudicators, all of which changed from ABSENT under the rule-based adjudicator to UNRESOLVED under Sol. Seven of these nine cases had been correctly classified by the rule-based adjudicator. Sol corrected none of the two rule-based errors.

## Discussion

The central result was a separation between orchestration and adjudication. GPT-5.6 Sol reached the same final classifications as fixed and exhaustive control while requesting fewer follow-up analyses but performed substantially worse when given final authority over the identical evidence record. The loss in performance cannot be attributed to missing tools or different data acquisition because measurement, available evidence and evidence state were held constant in the role-swap comparison. It arose from how the same evidence was converted into a biological decision. This distinction is consistent with broader calls to evaluate scientific agents at the level of individual capabilities rather than treating end-to-end performance as a single property (Chen et al., 2025; Liu et al., 2026).

The controller result clarifies what the LLM contributed. Sol did not discover measurements unavailable to the deterministic system; it selected a shorter analytical route to the same endpoint. Tool-augmented agents have repeatedly shown value in this role. ChemCrow improved scientific task performance by using an LLM to select and coordinate specialist chemistry tools rather than relying on the language model itself for exact chemical operations (Bran et al., 2024). GeneAgent similarly improved gene-set interpretation by retrieving evidence from curated biological databases and using that information to verify model-generated claims (Wang et al., 2025). In single-cell analysis, planning, retrieval and self-reflection contributed differently to task performance, further showing that agent value can arise from how analyses are organized rather than from a single end-to-end reasoning capability (Liu et al., 2026).

Biological adjudication is c different from tool selection from a computational context. An autoregressive language model is trained to estimate the probability of a token given preceding context, *P*(*x*_*t*_ ∣ *x*_<*t*_), whereas biological adjudication more closely resembles discrimination among competing hypotheses conditional on measured evidence and biological context, conceptually *P*(*H*, ∣ *EC*)(Brown et al., 2020). The is important because biological evidence is rarely self-interpreting. Sequence similarity can support common ancestry without establishing identical function, and functional transfer becomes particularly difficult when orthologues, paralogues and related protein families share overlapping sequence or domain evidence (Lee et al., 2007). Genomic machine-learning studies likewise show that dependence structure, confounding and dataset composition can change the meaning and apparent predictive value of biological features (Whalen et al., 2022). An LLM can combine such observations fluently, but its language-model objective does not itself provide an explicit, calibrated biological decision rule. Calibration can also deteriorate when language models are transferred to new tasks, even when they exhibit useful self-evaluation on familiar settings (Kadavath et al., 2022).

This distinction has begun to appear empirically in biological agent studies. Zhang (2026) supplied different frontier agents with the same deterministic protein evidence and found that they reliably retrieved and organized the input but produced only moderately reproducible biological relevance assignments; skill augmentation improved evidence handling and traceability without eliminating biological overinterpretation. More recently, Kim and Romero (2026) found that frontier agents generally selected appropriate tools, whereas much of the remaining performance gap arose from shallow evaluation of candidates, insufficient comparison of alternatives and premature termination. These observations are consistent with the present role-swap experiment, which supports that selecting an appropriate analysis and deciding whether the resulting evidence is sufficient for a biological conclusion are separable capabilities.

The results also question whether routine workflow control requires a frontier general-purpose LLM. In the present experiment, the constrained non-LLM Jev controller reproduced the endpoint calls and analytical workload of Sol while operating at substantially lower controller latency and cost. The conclusion should remain task-specific, but it shows that model complexity should be justified by measurable gains rather than assumed to improve an agent merely because more general reasoning is available. Similar conclusions have emerged outside biology, where simpler agentless pipelines have matched or exceeded substantially more complex agent architectures at lower cost, and benchmark analyses have emphasized evaluating accuracy jointly with cost and complexity (Xia et al., 2024; Kapoor et al., 2024).

Jev is computationally better suited to the control layer than to biological adjudication. As a typed, non-LLM classifier, it maps a structured evidence state onto a constrained action space, which is exactly the problem faced by an agent router or tool caller. In this setting, Jev matched Sol’s controller behaviour and final endpoints while operating at much lower latency and cost. Its weakness appeared when the task changed from action selection to biological interpretation: when asked to adjudicate identical evidence, it tended to abstain rather than resolve competing biological hypotheses. This suggests a useful division of labour: compact classifiers may be preferable for routine routing and tool selection, while biological adjudication requires an explicitly validated decision layer rather than simply a faster controller.

The failure mode of the LLM adjudicator showed that Sol did not primarily replace correct negative calls with false positive calls; it converted nine ABSENT calls to UNRESOLVED, including seven true negatives, while correcting neither of the two errors made by the rule-based adjudicator. Abstention is not intrinsically undesirable. Selective-classification theory explicitly permits a model to reject difficult cases in exchange for lower error among the cases it accepts, producing a risk–coverage trade-off (El-Yaniv and Wiener, 2010; Geifman and El-Yaniv, 2017). However, abstention adds value only when uncertainty is concentrated in genuinely difficult or error-prone cases. Here, the additional abstention flagged both missed positives as uncertain but resolved neither, and would instead have referred seven cases already classified correctly by the explicit rules.

In biological analyses, demonstrating that a target is absent requires more than failing to observe strong positive evidence; plausible competing explanations must also be excluded. In gene annotation, related protein families can share substantial sequence or structural features, and similarity alone is therefore insufficient for specific functional assignment (Lee et al., 2007). Mature specialist systems address this problem by encoding family-specific evidence criteria rather than relying on unconstrained interpretation. The present findings suggest that explicit biological decision boundaries remain useful precisely where several individually plausible signals must be translated into a reproducible endpoint.

Taken together, the results favour a tiered architecture for scientific agents. Deterministic software should generate quantitative measurements; a constrained controller should handle routine routing when the action space is well defined; a more capable LLM can be reserved for cases requiring open-ended selection among competing analyses; and final high-consequence biological calls should be grounded in explicit, testable decision criteria or referred for expert review when those criteria cannot resolve the evidence. This architecture is consistent with GeneAgent’s use of curated databases for external verification, Zhang’s recommendation for deterministic evidence tables and explicit scoring, and ScienceAgentBench’s argument for evaluating individual components before claiming end-to-end scientific autonomy (Wang et al., 2025; Zhang, 2026; Chen et al., 2025).

The results should not be interpreted as LLMs are generally unsuitable for biological interpretation. The experiment involved one gene-family endpoint, one frontier LLM configuration and a balanced cohort of 20 genomes. Multimodal and biology-specific models may ultimately improve biological adjudication by learning directly from sequence, structure, ontology and interaction data rather than relying predominantly on natural-language representations; emerging protein-function models are already exploring such architectures (Simon et al., 2024; Fallahpour et al., 2026). The present result demonstrates that competence in workflow orchestration cannot be assumed to imply competence in final biological adjudication, and those roles should therefore be evaluated separately.

In this study, the LLM added value by determining which analysis was worth performing next. It did not improve the final interpretation of the evidence it helped acquire. Scientific-agent design should therefore treat orchestration and adjudication as separate computational problems, assign each to the mechanism that performs it most reliably, and use abstention as a calibrated route to further analysis or human review rather than as a substitute for a biological decision.

## Data availability

All data underlying the reported results, including the prospective case manifest, locked truth labels, prediction records, evidence-state hashes, controller outputs, scoring tables and provenance manifests, are included in the archived study release at Zenodo (https://doi.org/10.5281/zenodo.22917959). The bacterial genome assemblies analysed in this study are publicly available from NCBI RefSeq; accession numbers and SHA-256 checksums required to identify the exact assemblies used are provided in the archived release. No new sequencing data were generated in this study.

## Code availability

Code and frozen study artifacts required to reproduce the reported analyses are available in the Genome Skeptic reproducibility release, archived at Zenodo: https://doi.org/10.5281/zenodo.22917959.

## Author contributions

A.S. conceived the study, developed the methodology and software, performed the analyses, prepared the figures and wrote the manuscript.

